# A scalable analytical approach from bacterial genomes to epidemiology

**DOI:** 10.1101/2021.11.19.469232

**Authors:** Xavier Didelot, Julian Parkhill

## Abstract

Recent years have seen a remarkable increase in the practicality of sequencing whole genomes from large numbers of bacterial isolates. The availability of this data has huge potential to deliver new insights into the evolution and epidemiology of bacterial pathogens, but the scalability of the analytical methodology has been lagging behind that of the sequencing technology. Here we present a step-by-step approach for such large-scale genomic epidemiology analyses, from bacterial genomes to epidemiological interpretations. A central component of this approach is the dated phylogeny, which is a phylogenetic tree with branch lengths measured in units of time. The construction of dated phylogenies from bacterial genomic data needs to account for the disruptive effect of recombination on phylogenetic relationships, and we describe how this can be achieved. Dated phylogenies can then be used to perform fine-scale or large-scale epidemiological analyses, depending on the proportion of cases for which genomes are available. A key feature of this approach is computational scalability, and in particular the ability to process hundreds or thousands of genomes within a matter of hours. This is a clear advantage of the step-by-step approach described here. We discuss other advantages and disadvantages of the approach, as well as potential improvements and avenues for future research.

## 1. Introduction

Over the past decade, the cost and time required to sequence whole bacterial genomes has reduced dramatically [1]. Sequencing is frequently applied to many or all isolates in local outbreaks, or to a high proportion of cases in more endemic situations, as well as large retrospective and longitudinal collections. This genomic data has huge potential to deliver new insights into the evolution and epidemiology of bacterial pathogens, which can lead to better control measures. However, the lack of scalable methodology for analysis of this genomic data represents an important bottleneck for the realisation of their full potential.

A gold standard for the analysis of pathogen genomic data has been set by the integrated phylogenetic frameworks implemented for example in BEAST [2] and BEAST2 [3]. These phylodynamic tools were originally conceived for viral genetics and are still mostly used for that purpose, but have also been increasingly applied to bacterial genomic data [4]. One of the strengths of these tools is that they can infer a dated phylogeny by combining the genomic data with the dates of isolation, resulting in estimates for the dates of the common ancestors in the phylogeny. Such dated phylogenies are extremely useful to draw epidemiological interpretations from the genomic data, as we will see. Another advantage of the integrated phylogenetic frameworks is that they include a number of powerful extensions, for example to use relaxed clock models [5], to estimate past population dynamics [6], geographical spread [7–9] or transmission between hosts [10,11]. This integrated approach has many natural advantages but also limitations especially in terms of scalability to analyse larger datasets.

These limitations of the integrated approach are especially important in bacterial genomics, where the genomes are orders of magnitude longer than in viral genetics and often subject to recombination. The ClonalOrigin model [12] of bacterial evolution has been integrated into BEAST2 [13], but the resulting algorithm is too computationally intense to be applied to whole genome datasets. Here we present an alternative step-by-step approach.

The step-by-step approach is illustrated in Figure 1. In the first step, a phylogeny is constructed from a genomic alignment in a way that accounts for recombination events. In the second step, this phylogeny is dated. In the third step, the dated phylogeny is interpreted in terms of a number of epidemiological properties. Many software packages are available to perform each of these steps, including but not limited to the ones named in Figure 1, although it is worth noting that many of these tools have emerged only in the past few years, and so are still work in progress and expected to improve in the near future. In this article we review each of the steps of this approach in turn. We also pay special attention to the ‘cracks’ between the steps, since these are often ignored in articles that focus on each of the steps rather than the whole step-by-step approach. Finally, we demonstrate the usability of this approach by applying it to a complete collection of *Staphylococcus aureus* ST239 genomes.

**Figure 1:**
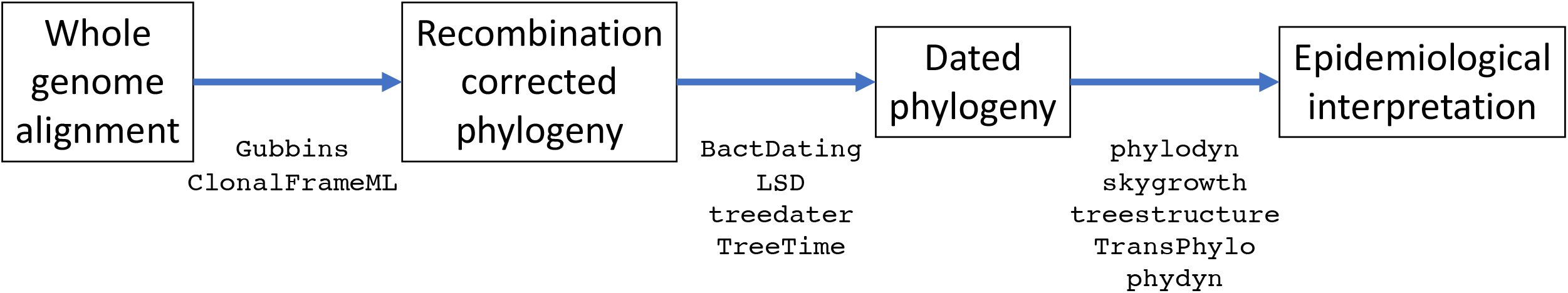
Overview of the step-by-step analytical approach. The names of some of the software tools that can be used in each step are indicated under the arrows.

## 2. Recombination-aware phylogenetic analysis

Even a relatively low amount of recombination can invalidate the results of phylogenetic tools if not accounted for [14,15]. It is therefore essential to detect recombination events to correctly reconstruct the clonal genealogy, that is the phylogenetic relationship between genomes when the ancestral lines of recipient cells rather than donor cells are followed for each ancestral recombination events. Special phylogenetic methods have been developed for this purpose, including Gubbins [16] and ClonalFrameML [17] which is based on the ClonalFrame model [18]. However, these tools are often underexploited, typically to build a recombination-corrected tree without paying attention to the recombination events and regions that have been detected.

A lot can be learnt from studying the inferred recombination events themselves. Recombination is useful to help us understand how species are being formed [19] and the population structure within species, especially when the origin of recombination events is being investigated [20]. These recombination patterns often reflect important driving evolutionary forces such as ecology [21], adaptation [22] or selective pressures [23]. For example in *Streptococcus pneumoniae*, recombination events have been shown to be driven by antibiotic usage in a localised dataset [24] and by immune pressure in a global collection of the PMEN1 lineage [25]. The latter study also represents a good example of how the temporal signal can become much clearer once recombination is correctly accounted for [25,26]. Recombination is also useful for the analysis of genome-wide associations between genotypes and phenotypes, since it separates new genetic variants from their original genomic background [27].

Accounting for recombination when reconstructing phylogenies is an important starting point for many epidemiological studies. A method often used is to extract from the genomic alignment the sites that have not been affected by recombination and to build a phylogeny using these sites only. Both Gubbins and ClonalFrameML are often used in this way, to create a recombination-free alignment which is then passed on to BEAST. However, this method works only if relatively few recombination events happened throughout the tree. For example, consider the simulated dataset shown in Figure 2. The true clonal genealogy is shown in Figure 2A and the true recombination events that happened on each of the branches are shown in Figure 2B. These data were simulated using a standard coalescent model for the phylogeny [28], a strict clock model of mutation with rate θ/2=0.005 per site, a model of recombination coming from external sources [18] with initiation rate ϱ/2=0.001 per site, average length of recombination *δ*=1500bp and distance of the source ν=0.05. For clarity we used a relatively small dataset of 20 sequences of 100,000bp each. In this simulated dataset, there was not a single site that was not affected by recombination on at least one of the branches. On the other hand, every branch had some sites unaffected by recombination (Figure 2B).

**Figure 2:**
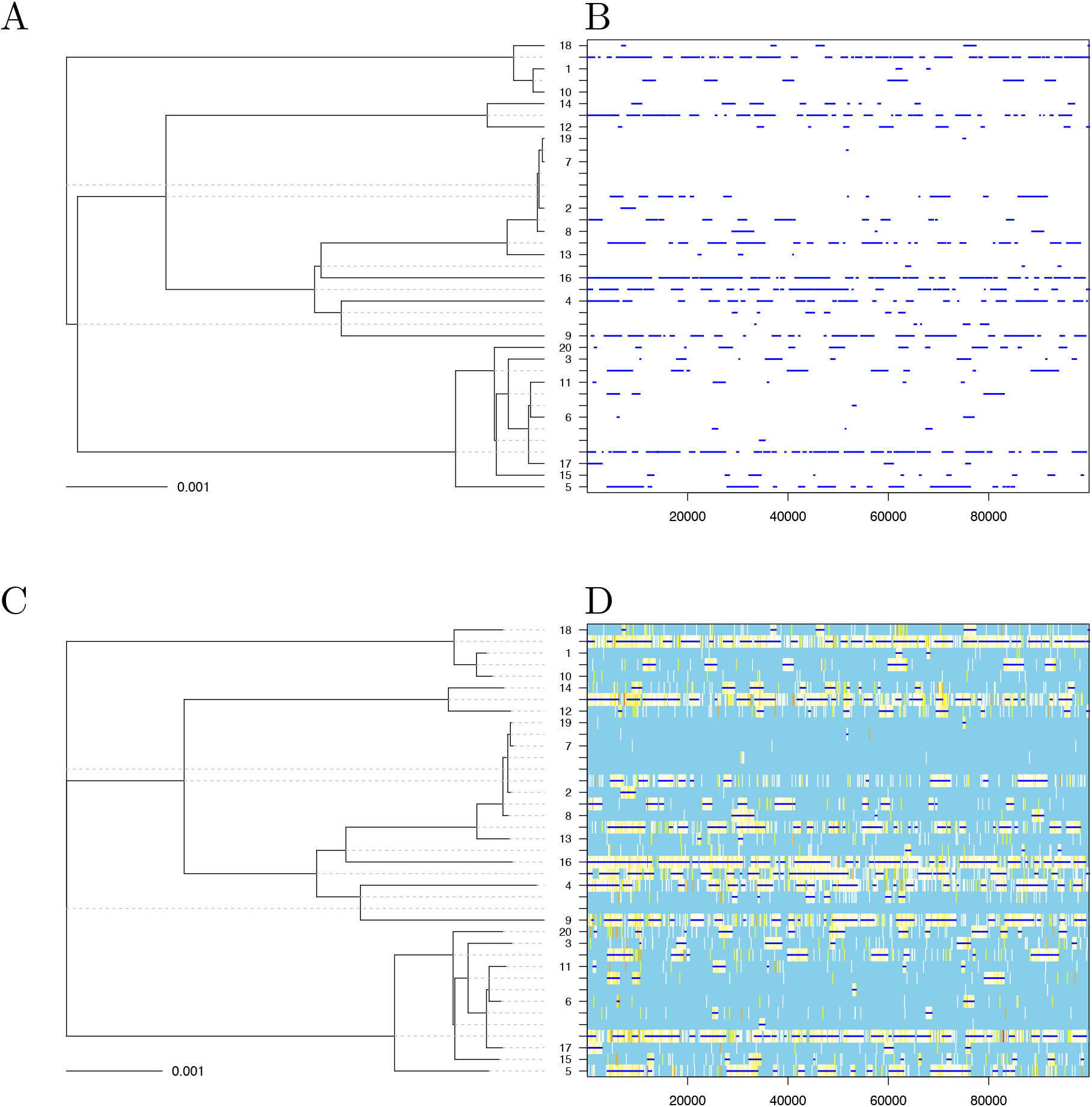
Illustration of the effect of recombination on phylogenetic inference. A phylogeny was simulated (A) with recombination events happening on the branches at a constant rate (B). ClonalFrameML was applied to this simulated dataset, resulting in a good reconstruction of both the clonal genealogy (C) and recombination events (D).

We applied ClonalFrameML [17] to this dataset using a PhyML tree [29] as starting point. The reconstructed clonal genealogy is shown in Figure 2C and the inferred recombination events are shown in Figure 2D, and they are in very good agreement with the true simulated tree and events shown in Figures 2A and 2B. ClonalFrameML correctly inferred that there was not a single site unaffected by recombination on at least one of the branches. Therefore an alignment containing only the non-recombinant sites would contain no sites, and could not be used as a starting point for further analysis. On the other hand, the inferred clonal genealogy shown in Figure 2C can be used in our proposed step-by-step approach. It has the same topology as the true clonal genealogy (Figure 2A) and very similar branch lengths, with a weighted Robinson-Foulds distance [30] of 0.005 between the true and ClonalFrameML trees. Gubbins [16] was also applied to this dataset using RAxML [31] as a tree builder. The correct topology was inferred, with a weighted Robinson-Foulds distance of 0.03 between the true and Gubbins trees.

## 3. Dating the ancestors in a phylogeny

Once a recombination-corrected tree has been reconstructed, it is possible to study the temporal signal in this tree and to date the common ancestors in the tree. Multiple software tools have recently been developed to perform dating on a phylogeny, including BactDating [26] which is specifically aimed at bacterial genomes, but also LSD [32], treedater [33] and TreeTime [34]. BactDating uses Bayesian statistics, whereas treedater and TreeTime are based on maximum likelihood, which is identical to a Bayesian maximum a-posteriori (MAP) approach assuming a uniform prior on dates as previously proposed [35]. It is often important to use a relaxed clock model in this step that allows the evolutionary rate to vary between lineages [5]. An additive relaxed clock model has recently been developed which is more biologically realistic and leads to better dating of pathogen phylogenies than previous relaxed clock model [36].

In our proposed step-by-step approach, the reconstruction of a dated phylogeny and its epidemiological interpretation are separated. One disadvantage of this is that the prior (or lack-of) on dates used to reconstruct the dated phylogeny is not the same as the one that would be implied by the epidemiological models used in subsequent analyses. This statistical issue could be resolved for example by considering the difference in tree distribution between the models used for the dating and the epidemiology and applying an importance sampler to correct for this difference [37]. However, this difference is often small enough to be ignored in practice, especially if the method used to build the dated phylogeny was based on the likelihood only, or if a mild prior was used such as the coalescent with constant population size [28].

To illustrate this, we simulated five years of an outbreak model [38] with within-host diversity N_e_g=0.25 year, basic reproduction number R_0_=2, generation time distribution Exponential(1) in years and sampling proportion *π*=0.1. A total of 59 cases were sampled in this outbreak, with the samples being related as shown in Figure 3A. We applied a strict clock model to this dated phylogeny with a rate μ=5 substitutions per year which is of the same order of magnitude as many bacterial pathogens [39]. This undated phylogeny was then used, along with the known dates of sampling, to infer a dated phylogeny using BactDating [26] with prior set to the coalescent with constant population size [28]. This prior is very different from the outbreak model that was used to generate the phylogeny [38], which is not coalescent due to the host structure and where the population size is clearly growing since the reproduction number was greater than one. Figure 3B shows the inferred dating for this tree, which is in good agreement with the correct dates from the top part despite the complete difference between the epidemic model used for simulation and the coalescent model used for inference.

**Figure 3:**
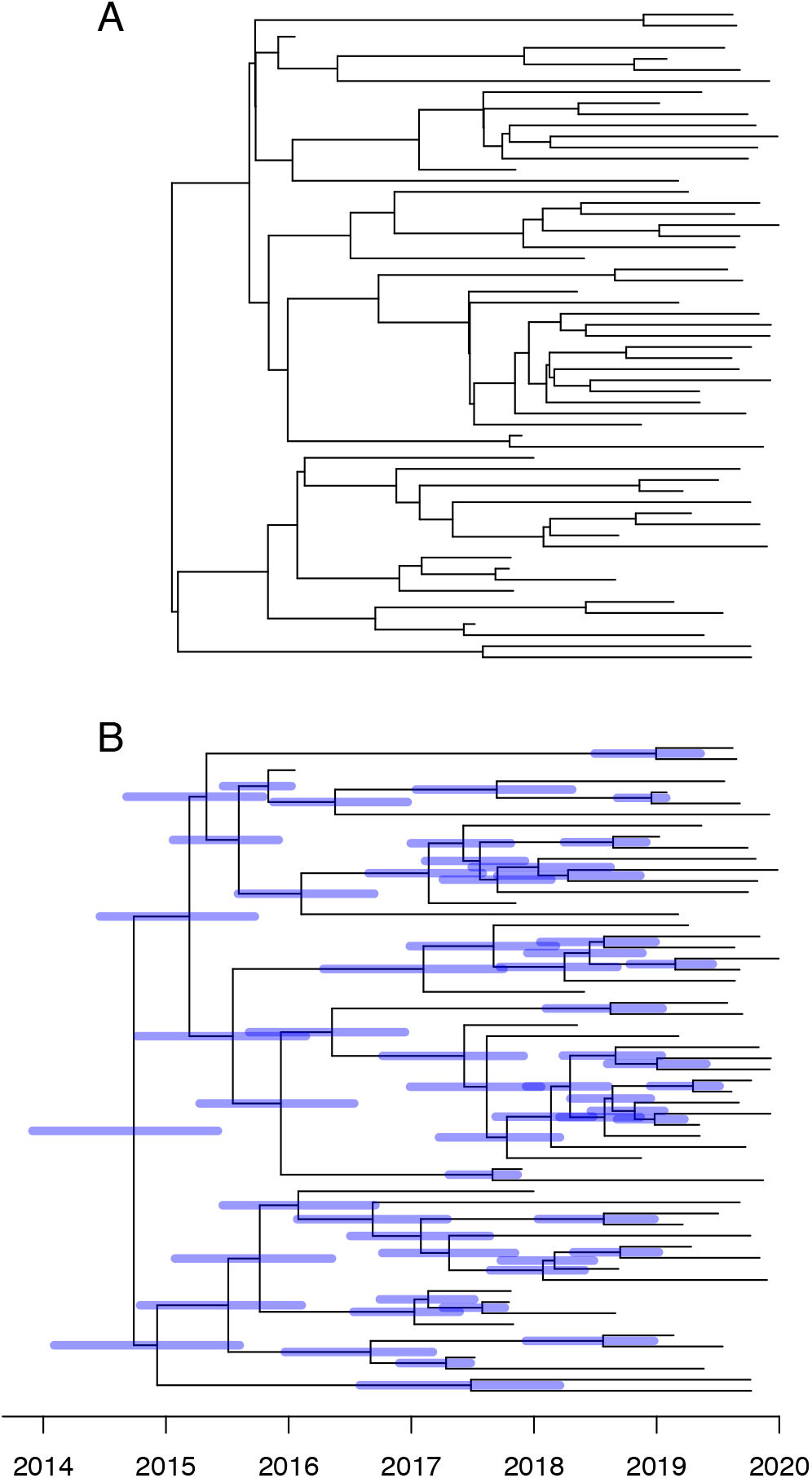
Illustration of the relative lack of effect of the prior model used for the inference of dated phylogeny. A dated phylogeny (A) was simulated from an epidemic model and dating was inferred (B) based on a coalescent model with constant population size.

At the same time as dating is performed, the substitution rate is typically estimated which provides a useful value to compare with previous estimates [39] in order to make sure that the dating is working as expected. Statistical methods can also be used to ensure that the temporal signal is significant, for example by comparing the fit of the data when the correct sampling dates are used against when all the dates are forced equal [40], or using a permutation test on the sample dates [41]. These methods require to perform several runs of the dating method, and it is therefore useful for this to be as fast as possible, which is achieved in our step-by-step method by separating the phylogenetic inference from the dating.

Furthermore, the root of the phylogeny is typically estimated during the dating step, since the trees generated by standard phylogenetic tools are not rooted whereas dated trees are always rooted by definition, with the date of the root being the date of the last common ancestor of the whole sample. If the root has already been determined robustly, for example using one or ideally several closely related outgroups [42], then this information can be preserved during the dating. If on the other hand the root is undetermined, or arbitrarily selected for example using the midpoint method [43], then the fact that dating the phylogeny simultaneously performs rooting provides an additional reason for dating the tree, which becomes much more informative in terms of epidemiology once it is dated and rooted.

## 4. From dated phylogeny to epidemiology

A dated phylogeny is very useful to learn about the epidemiology of the bacteria under study, and sometimes the dating directly provides answers to questions of interest beyond the age of pathogens [44]. For example, several antibiotic resistant lineages have been dated to have emerged around the time when the corresponding antibiotics were started to be used, highlighting the link between consumption and resistance [45,46]. As another example, the dating of the common ancestors between pairs of *Clostridium difficile* patients in a hospital allowed to rule out transmission for many pairs and to conclude that nosocomial transmission was less frequent than previous thought [47].

It can often be useful to identify clusters of significantly similar genomes in a dataset. The most commonly used approach is to use a separate dedicated algorithm that uses the genomic data for this purpose, such as HierBAPS [48], fastbaps [49] or PopPunk [50], and overlay the results of this clustering analysis onto the phylogeny using colours for example. Another approach is to use additional non-genomic data to do the clustering given the phylogeny, as performed for example by AdaptML [51], treebreaker [52] and treeSeg [53]. Finally a third option is to try and identify directly on the dated phylogeny the lineages that seem to be ruled by different dynamics, for example using treestructure which does not rely on an explicit phylodynamic model [54] or CaveDive [55] which is focused on the detection of clonal expansions.

The dated phylogeny can also be used as a starting point for further analysis. In particular, past variations in the bacterial population size have a direct effect on the shape of the dated phylogeny, so that the population size through time can be estimated and presented as a skyline plot [56]. The methodology for performing such an analysis was originally developed within BEAST which simultaneously estimates the dated phylogeny [6], but for our step-by-step approach we need to estimate the demographic function from a dated phylogeny, and several software tools have recently been released for this purpose including phylodyn [57,58], skygrowth [59] and mlesky [60]. Beyond a simple model of varying population size, it is also possible to fit an epidemiological compartmental model such as the susceptible-infected-recovered model [61], and therefore to estimate the parameters of this model such as the transmission rate or removal rate. Such an inference can be achieved by formulating a structured coalescent model that corresponds to the compartmental model [62,63]. Existing software for fitting such a model to a given dated phylogeny include rcolgem [64] and phydyn [65]. The same methods based on the structured coalescent can also be applied to a dated phylogeny in order to reconstruct past geographical migrations [9], although such phylogeographic inference is much more often based on discrete trait analysis, for example using the ace command from the R package ape [66] or in the NextStrain platform [67]. The worldwide spread of the current pandemic of *Vibrio cholerae* has been described using such techniques [68,69].

When the genomes are densely sampled within an epidemic, it can be useful to try and reconstruct the transmission tree of who infected whom [70]. Within-host diversity and evolution is significant for many bacterial pathogens which blurs the relationships between transmission tree and phylogeny [71]. However, TransPhylo can infer the transmission tree from a dated phylogeny in a way that accounts for within-host evolution [38,72,73]. Significant uncertainty typically remains in the inferred transmission tree, which is captured by the use of Bayesian statistics within TransPhylo. More precise inference can sometimes be obtained by combining the genomic inference with epidemiological data [74].

A drawback of separating the dating step from the interpretation step is that the uncertainty in dating is typically not passed on to the epidemiological analysis. This can be achieved by running on multiple samples from the posterior of dated phylogeny and averaging the results [75], or reweighting according to the posterior probability in the epidemiological analysis [37], but in practice the phylogenetic uncertainty is usually not accounted for. However, this is not often a significant issue in practice. To illustrate this, we simulated a dataset for a small outbreak with just ten cases, using an epidemic model [38] with basic reproduction number R_0_=1, within-host diversity N_e_g=0.25 year, mean generation time of 1 year, sampling proportion of *π*=0.5 and a strict clock model with rate μ=5 substitutions per year. The dated phylogeny was inferred using BactDating [26] and we extracted the first (after burnin) and the last trees sampled by the MCMC, as shown in Figures 4A and 4C. We then reconstructed the transmission events using TransPhylo [38] separately for each of these two dated trees, as shown in Figures 4B and 4D. In spite of small differences in the two dated phylogenies, the inferred results in terms of transmission chains were very similar.

**Figure 4:**
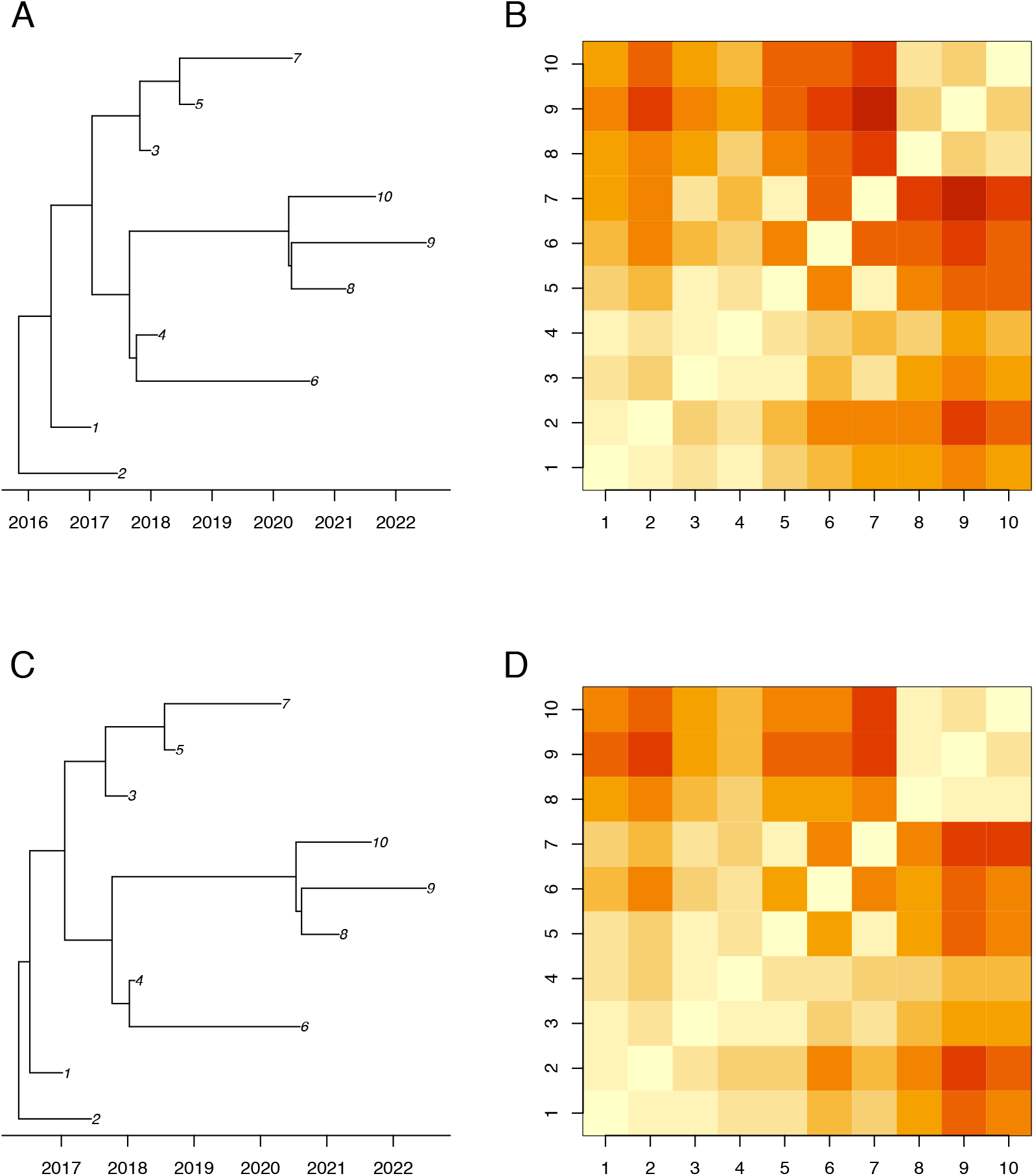
Illustration of the relative lack of effect of the uncertainty in the reconstructed dated phylogeny on interpretation as a tree transmission trees. Two dated phylogenies were sampled from the posterior (A and C) and a separate inference of the transmission tree was performed for each one (B and D). The coloured matrices represent the distance between pairs of cases in number of transmission links, with red coding for direct transmission and yellow coding for a distance of ten links.

## 5. Example of application

To illustrate the use of the step-by-step approach from bacterial genomes to epidemiology, we apply it to a state-of-the-art dataset, using only a standard laptop computer and paying particular attention to the time taken by each step. We collected all available genomes of *Staphylococcus aureus* ST239 (Table S1). This collection is made of 521 assembled genomes, only small subsets of which had been comparatively analysed in previous studies [76–79]. The genomes were collected between 1982 and 2010 from all parts of the world (451 from Asia, 46 from Europe, 18 from Americas, 2 from Africa, 2 from Oceania and 2 unknown). All genomes were aligned using MuMMER v3.1 [80] against the reference genome TW20 which is a member of ST239 [81] and therefore included in the collection. This resulted in a reference-anchored alignment that took only a few minutes to generate, since each pairwise alignment against the reference genome can be performed in parallel. Alternatively, assembly pipelines are often based on reference-based mapping of the sequencing reads, for example using BWA [82] and SamTools [83]. This can also be performed in parallel and results in a similar reference-anchored alignment.

A first phylogeny was built using PhyML v3.3 [29] which took approximately 3 hours. This was used as the starting point to build a recombination-corrected phylogeny using ClonalFrameML v1.12 [17], which took approximately two days to run. The same analysis using Gubbins v2.4.1 [16] gave very similar results, and took approximately one day to run. This step currently represents a clear bottleneck in the application of the step-by-step approach, which should be addressed in the near future through the development of new parallelised algorithms. Significant recombination was found, with a total of 198 recombination events detected throughout the phylogeny. The relative rate of recombination versus mutation was estimated to be R/θ=0.144, meaning that on average mutation events were about 7 times more frequent than recombination events. The mean length of recombination events was estimated to be *δ*=619 bp which is in good agreement with previous estimates for S. aureus [17,84,85]. The mean distance between donor and recipient was estimated to be ν=0.31%, which corresponds approximately to the distance between ST239 and some of its closest relatives such as CC8 [86]. The relative effect of recombination versus mutation was therefore estimated to be r/m = R/θ **×** ν **×** *δ* = 0.28, so that 3 to 4 times more substitutions are caused by mutation than by recombination. These results confirm that recombination plays a role in *S. aureus* evolution, although not as dramatic as in some other bacterial pathogens [14,87,88].

We detected a strong temporal signal in the recombination-corrected phylogeny on the basis of a regression analysis of root-to-tip distances against isolation dates (R^2^=0.57, p<10^−4^). We therefore computed a dated phylogeny using BactDating v1.1 [26] under the additive relaxed clock model [36]. This step took approximately 3 hours to run for 10^6^ MCMC iterations, and the inferred dated phylogeny is shown in Figure 5A. The isolation dates were unknown for 36 of the 521 genomes (Table S1), but BactDating can accommodate this. The evolutionary rate was estimated to be 7.05 substitutions per year throughout the genome, with credible interval between 6.43 and 7.67. This estimate is in good agreement with several previous estimates in ST239 [76,78] and other lineages of *S. aureus* [46]. The root of the ST239 was estimated to have existed in 1958, with credible interval ranging between 1951 and 1965. This is again in good agreement with previous estimates and coincides with penicillins being increasingly used to treat bacterial infections [76,78,89].

**Figure 5:**
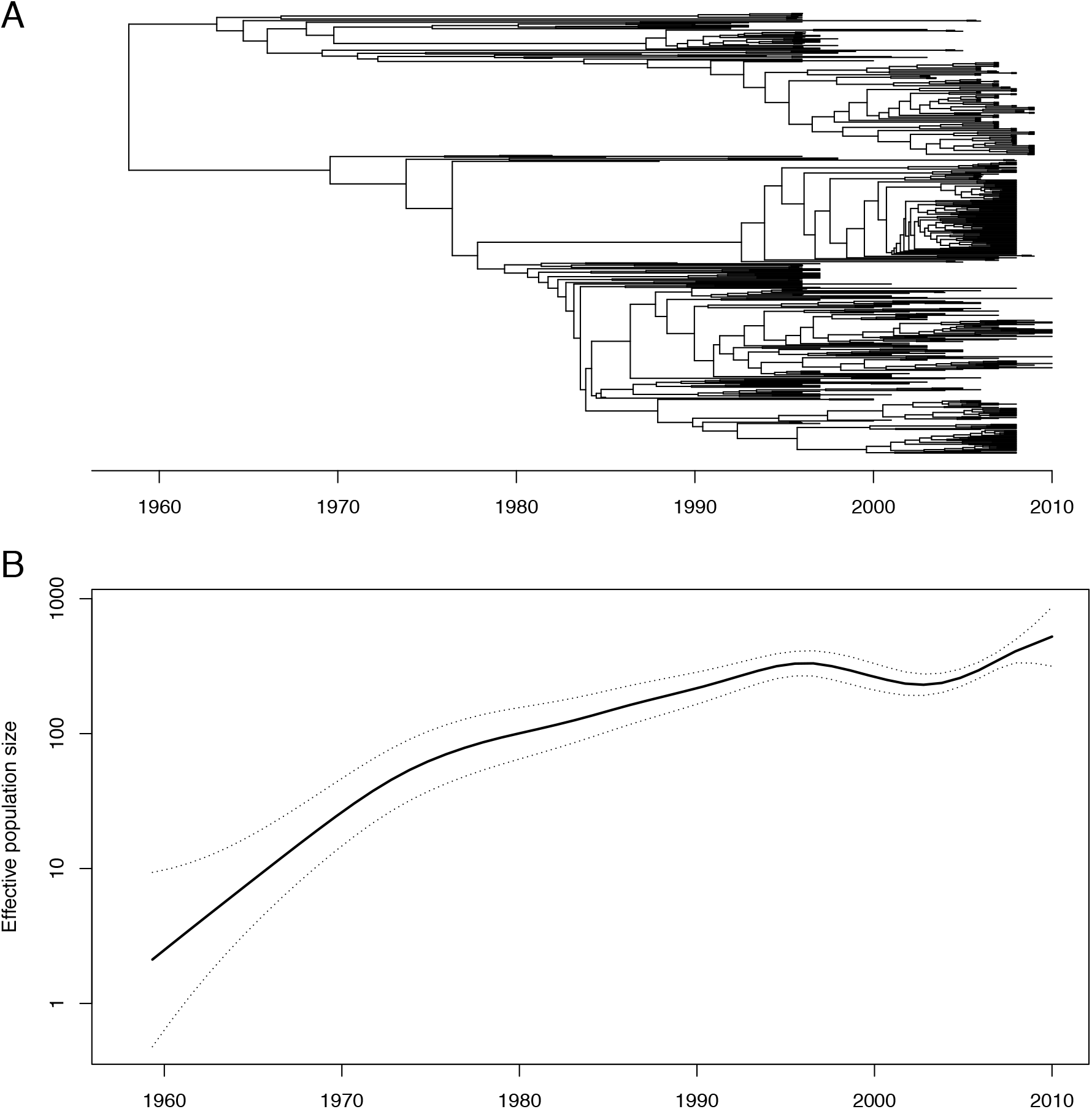
Example of application of the approach to a collection of *Staphylococcus aureus* ST239 genomes. The dated phylogeny was inferred using BactDating (A) and the past population size dynamics was inferred using skygrowth (B).

We used the dated phylogeny as input into treestructure v0.1.2 [54] to determine whether there were significant differences in the phylodynamic properties of sublineages within the tree. This analysis took less than a minute to perform, and found no significant differences, which means that the whole tree can be treated as a whole in phylodynamic reconstructions [54]. We therefore applied skygrowth v0.3.1 [59] to the whole dated tree using the maximum a-posteriori method. This analysis took less than a minute and the estimated demographic function is shown in Figure 5B, with an approximately exponential rise of the effective population size between 1960 and 1995, and a plateau between 1995 and 2010. This is in good agreement with previous skyline analyses of ST239 [78,89]. We do not seek to say more about the epidemiological dynamics of ST239 since our aim with this application was to test the applicability of the step-by-step method to a relatively large dataset, rather than study it in detail.

## 6. Discussion

The step-by-step approach has several drawbacks compared to an integrated approach. A practical disadvantage is that multiple tools need to be applied one after the other, with the need to make sure that the output of one tool is a suitable input for the next tool. The software tools have been developed separately, and format conversion is sometimes required when combining them, which introduces a risk of error being made. Method developers should make every effort to minimize this risk, for example by providing practical examples of source code combining new tools with pre-existing ones, and including verifications in each tool that the input is formatted as expected.

Another concern with the step-by-step approach relates to statistical soundness. In an integrated approach, a complex model is formed by combining multiple simpler models into a consistent whole, for example a model describing how the pathogen population size varied over time, another model describing how these fluctuations affect the genealogy and yet another model describing how mutation and recombination events affect the genomes given the genealogy. Inference is then performed on the combined model, with all uncertainties being accounted for simultaneously and in all directions: for example the uncertainty on a mutation event will feed into the uncertainty on the past population size, and vice-versa. By contrast, in the step-by-step approach, each of the tool makes separate modelling assumptions, which may not always by consistent with each other. An example of this was discussed in Section 3, where the prior used for the reconstruction of a dated phylogeny was not the correct one, but Figure 3 showed that the result can still be correct. Furthermore, the uncertainty can only be passed from one tool to the next in the order that they are being applied, and even in this direction it is frequent to use the best estimate from one method as the starting point of the next, without passing any uncertainty. Again this is not necessarily a problem in practice, as illustrated in Figure 4 where the uncertainty on the phylogeny had little effect on the uncertainty of the transmission tree. From a statistical point of view, the integrated approach therefore represents a gold standard, although statisticians have recently noted that joint inference under a combined model carries the risk that misspecification in any of the model parts can affect estimates from the others in unpredictable ways [90]. Further research is needed on this in the context of genomic epidemiology, as well as research on how to avoid the statistical issues described above with the step-by-step approach.

A key advantage to the step-by-step approach we described is that by breaking down the problem into simple steps, it becomes easier to solve, a strategy often called “divide and conquer” in the computer science literature. The running time is greatly improved compared to an integrated approach, which quickly becomes intractable as more model components are combined into a large model. An example of this concerns the difficulty to integrate recombination into a phylodynamic framework [13]. A similar situation occurs when aligning sequences and building a phylogeny: in principle alignment and phylogeny would benefit from being performed simultaneously [91,92] but in practice this is too computationally challenging. The lower running time of the step-by-step approach also means that it is more scalable to the large numbers of bacterial genomes currently available, and this scalability is probably the main reason for a recent increase in popularity [67,93].

Perhaps even more importantly, a counterintuitive advantage of the step-by-step approach is that it is less automatic than the integrated approach. Although this may seem like a disadvantage, the fact that several software tools have to be applied one after the other brings great benefits. It allows the user to check after each step that the result makes sense before carrying the next step. For example, if a phylogeny is clearly wrong due to contamination during sequencing, there is no point trying to apply dating of the nodes or interpreting the phylogeny in terms of epidemiology. Since each tool is focused on a simpler task, it is easier for the user to check the validity of the assumptions made, and if needed to compare models or the results of several software tools, or apply more complex models since each step is relatively quick. These checks and refinements provide the user with a better understanding of their data and the analysis process, rather than relying on “black-box” or “turn-key” analysis. This is one of the most important advantages of the step-by-step approach, since it creates good conditions for a balanced interpretation of the data and results.

## Acknowledgments

We acknowledge funding from the National Institute for Health Research (NIHR) Health Protection Research Unit in Genomics and Enabling Data.

